# Single Cell App: An App for Single Cell RNA-sequencing Data Visualization, Comparison and Discovery

**DOI:** 10.1101/2021.10.09.463793

**Authors:** Mathew G. Lewsey, Changyu Yi, Oliver Berkowitz, Felipe Ayora, Maurice Bernado, James Whelan

## Abstract

The Single Cell App is a cloud-based application that allows visualisation and comparison of scRNA-seq data and is scalable according to use. Users upload their own or publicly available scRNA-seq datasets after pre-processing to be visualised using a web browser. The data can be viewed in two colour modes, Cluster - representing cell identity, and Values – level of expression, and data queried using keyword or gene identification number(s). Using the app to compare four different studies we determined that some genes frequently used as cell-type markers are in fact study specific. Phosphate transporter and hormone response genes were exemplary investigated with the app. This showed that the apparent cell specific expression of *PHO1;H3* differed between GFP-tagging and scRNA-seq studies. Some phosphate transporter genes were induced by protoplasting, they retained cell specificity, indicating that cell specific stress responses (i.e. protoplasting). Examination of the cell specificity of hormone response genes revealed that 132 hormone responsive genes display restricted expression and that the jasmonate response gene *TIFY8* is expressed in endodermal cells which differs from previous reports. It also appears that *JAZ* repressors have cell-type specific functions. These differences, identified using the Single Cell App, highlight the need for resources to enable researchers to find common and different patterns of cell specific gene expression. Thus, the Single Cell App enables researchers to form new hypothesis, perform comparative studies, allows for easy re-use of data for this emerging technology to provide novel avenues to crop improvement.

## Introduction

The ever-increasing throughput, declining cost per-sample and growing diversity of gene expression analysis methods has resulted in massive public repositories of freely accessible transcriptome data (Zhang et al., 2020). A variety of tools are available that provide user-friendly data access and interrogation to those plant biologists who are not experts in bioinformatics or primary data processing. However, these predominantly serve the most common and more established techniques, such as bulk RNA-seq or microarray analysis of whole organs. Single cell RNA sequencing (scRNA-seq) has emerged more recently as an extremely powerful technique to investigate the functions of individual cells and cell-types. Beyond visualisation, in recent years scRNA-seq has been applied to several plant species, generating valuable resources for the researchers to explore plant genomics at the single cell level (Denyer et al., 2019; Jean-Baptiste et al., 2019; Liu et al., 2021; Ryu et al., 2019; Xu et al., 2021; Zhang et al., 2021). However, these data are typically stored in a text-based format that must be processed and visualized for users to explore it. This creates a significant challenge for those plant scientists with limited computational experience, limiting data reuse.

Beyond visualisation there is a need for researchers to be able to compare different datasets of their choice, to design new experiments, to compare studies and to define cell specific genes using the parameters of their choice. While the publicly available RNA-seq and scRNA-seq repositories provide a valuable resource for a re-use of raw data, making them more accessible allows for their thorough investigation by experts in their field but with limited bioinformatics background. For RNA-seq data this is achieved by a variety of databases that have emerged in the last 20 years. The Bio-Analytic Resource for Plant Biology (BAR) and Genevestigator are notable examples of providing a repository and analytical tools to compare RNASeq data (Toufighi et al., 2005; Zimmermann et al., 2004). However, these visualisation tools are not compatible with highly dimensional and large data sets created by scRNA-seq, where expression of thousands of genes is measured in thousands of cells. This issue will only become more challenging as the cost will decrease over time. Furthermore, most plant biologists are not bioinformatics experts and are unlikely to be able to rapidly learn how to analyse scRNA-seq data with the currently available command line tools. This creates a significant obstacle to accessing and re-using public data. Moreover, substantial computing hardware is required to store and process scRNA-seq data, which costs tens to hundreds of thousands of dollars to acquire. When re-analysis is likely to be a short-term project, and not a continuous or ongoing requirement, it is difficult to justify such a substantial capital investment.

Tools have been developed to overcome these obstacles such as SC1 (Moussa and Mandoiu, 2021), Single Cell Explorer (Feng et al., 2019) and alona (Franzen and Bjorkegren, 2020). However, each of these has some disadvantages. For example, SC1 is not currently able to process plant scRNA-seq data. Single Cell Explorer needs to be installed on a Linux server by the users. Moreover, these tools primarily focus on scRNA-seq data processing whilst providing limited data visualization functions. Genevestigator has established a single-cell RNA-seq portal but is restricted to animal/human studies.

One of the central questions in single cell analyses is the identification of marker genes to determine cell types (Shaw et al., 2021). Stress responses diminish cell-specific signatures which impedes cell type assignment as shown for Arabidopsis roots (Jean-Baptiste et al., 2019). Similarly, in a study examining tissue specificity of gene expression in Arabidopsis leaves the tissue specific expression decreased with applications of chemicals that mimicked adverse growth conditions (i.e. oxidative stress) or hormones (Berkowitz et al., 2021). Hence it is difficult for non-expert researchers to test the robustness of marker genes defined by earlier studies, and to evaluate how consistent they may be under different conditions. Additionally, the increasing application of genomics to non-model systems means that the capability to define marker genes will be needed for a variety of plants species, as well as tools that allow researchers to compare markers between studies and species. PlantscRNAdb has compiled scRNA-seq datasets from plants and defined 26 326 marker genes of 128 different cell types from four plant species (Shaw *et al*., 2021). This is apparently high number shows the diversity and difference in approaches to defining marker genes. Researchers need to be able adjust the parameters to define marker genes to include new knowledge and allow discovery. Parameters set too strict or too lenient might mean that results are missed or meaningless.

To address this challenge we developed the Single Cell App. This tool focuses on interactive visualization of user-processed or public scRNA-seq data. All data storage and processing are conducted using a Microsoft Azure cloud-based platform, removing the necessity for users to purchase costly compute hardware. The app provides a web-based user-friendly graphical interface, making it appropriate for plant scientists who are not bioinformatics experts. Additionally, by providing these tools within an app, they can be rapidly and easily deployed and updated within a user’s institutional environment, greatly reducing challenges around installation and software maintenance. Overall, this cloud-based approach can scale readily dependent upon the needs of individuals or institutions. The Single Cell App can be obtained for local installation at (http://Single-Cell-Visualisation.loomesoftware.com).

## Results

### Overview of the single cell app

The Single Cell App is designed to provide an easily accessible environment for users new to scRNA-seq to explore data. It enables users to visualize cell expression profiles. The app is deployed within the user’s host institution IT infrastructure and once established, users connect over the internet using a web browser via a custom web address (URL) that can be associated with any institutional domain. This is achieved using the Loome Platform (https://www.loomesoftware.com/index) and Microsoft Active Directory.

There are three main screens in Single Cell App interface once the App is installed – Upload Experiment, Experiment Analysis and Experiment Status (Fig.1). Upload Experiment provides the functionality to upload gene expression data, clustering and gene ontology (GO) information and metadata associated with the data so that it can be identified as part of an experiment. Experiment Analysis provides the visualisation of the data for experiments that have been uploaded (Fig. 1), with rich controls to search, filter and modify the data that is being visualised. The user can also export the data being visualised in GO slim or comma-separated values (CSV) format.

**Figure 1.**
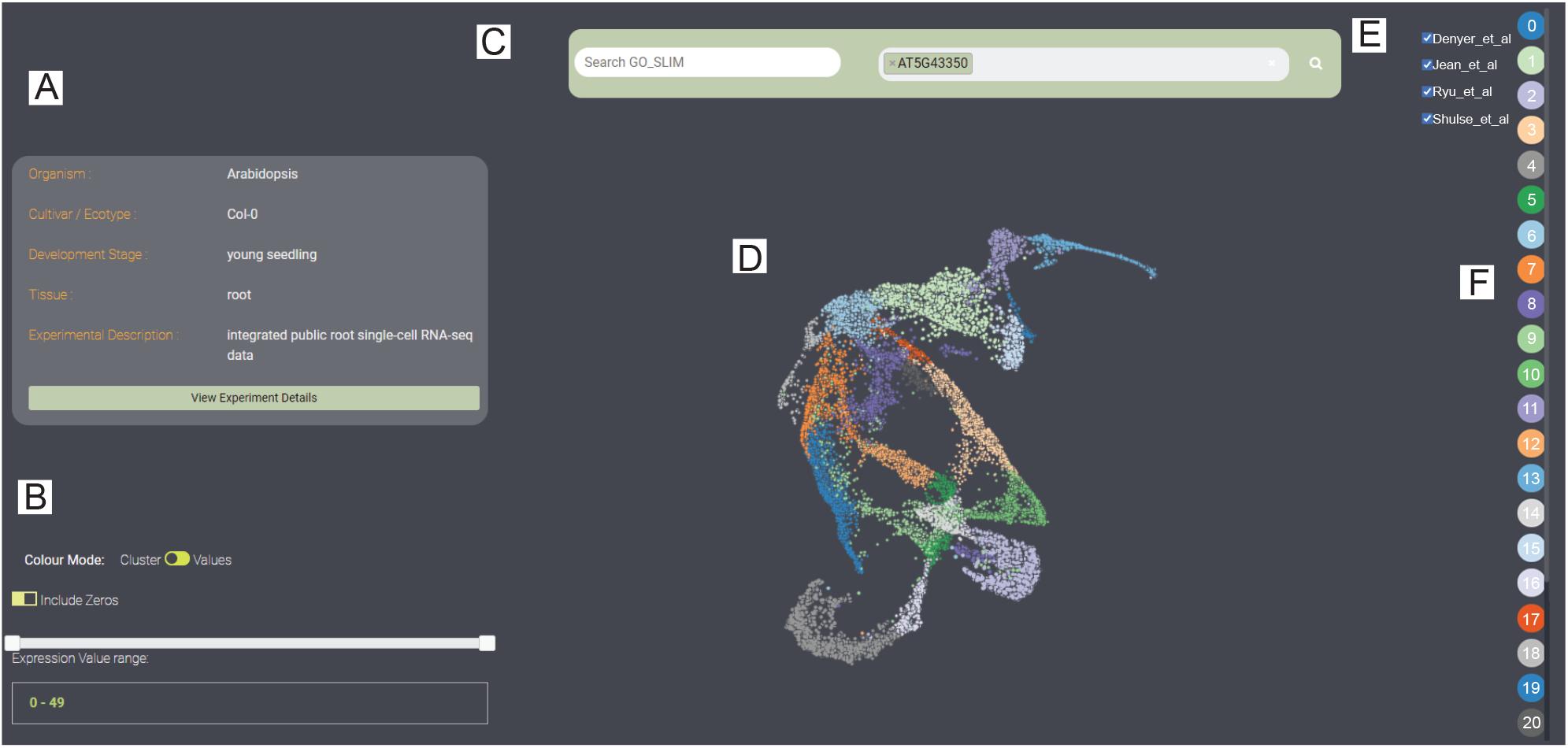
Screenshot of the Single Cell Browser Layout. (A) Meta information of the selected experiment. Clicking the “View Experiment Details” will provide more information about the experiment. (B) Color mode of displaying data and filtering cells based on gene expression level. In Cluster mode, each color represents a cell cluster while in Values mode, the color represents the expression level of a searched gene. Cells can be filtered by dragging the Expression Value range bar. (C) Search tab. Genes can be searched based on the Gene Ontology or gene id. (D) Three dimension of the UMAP clustering. Each dot represents a cell. (E) Check box for dataset selection, this can be used for showing cells from specified dataset. (F) Cluster legend. Each color represents a cluster, the number shows the cluster id. Clicking each cluster will hide/show the cells from this cluster.

### Utility of the single cell app

Users can upload experiments once the app has been installed on their own server by using the Upload Experiment function. Three data tables in CSV format are required to upload an experiment; 1) a gene expression matrix, 2) a meta information table describing each cell and 3) a GO slim file for the species of study. In the gene expression matrix, the first column is the gene ID and other columns are the unique cell barcode, the values represent the expression of each gene in each cell. The meta information table provides the three-dimensional coordinates from clustering analysis (e.g. T-SNE, UMAP), cell identity and other experimental information for each cell. The GO slim file contains the annotation of each gene. An example of these data tables has been provided in Supplemental Table 1-3. Notably, the system is species agnostic so long as all three files relate to the same species and use consistent nomenclature.

The main strength of our app is providing scRNA-seq data visualization with various interactive functions. Users can select different uploaded experiments from the drop-down list and examine the related meta information. The app provides a search function based on gene id and/or GO annotation, which allows users to explore and discover genes of their interest (Fig. 1C). For instance, one can search genes associated with phosphate-related biological processes by providing “phosphate” in the “Search GO_SLIM” box and then select an associated gene in the “Search GeneID” box. One can also search a gene of interest from the search box directly. Cells can be filtered based on the expression level of a searched gene, with cells above or below user-selected threshold masked. Moreover, the app provides two colour modes (Cluster and Values) to display the cells. In Cluster mode the colour represents cell identity (Cluster number) while in the Values mode colour indicates the expression level of cells (Fig. 1B). The app also provides a three-dimensional visualization of the cell clustering (Fig. 1D) and selection of cells based on datasets (Fig. 1E) or cluster identity (Fig. 1F).

### Characterising variation of four public Arabidopsis root scRNA-seq datasets

Roots provide an ideal tissue for analysing cell heterogeneity because high-quality ground-truth datasets exist in which all root cell types have been carefully identified, purified and analysed transcriptomically (Drapek et al., 2017; Shahan et al., 2021). Moreover, root development is very well characterised and roots are easily accessible and amenable to single-cell processing methods (Shahan *et al*., 2021). Several scRNA-seq studies have been performed in Arabidopsis root using the Columbia-0 (Col-0) genotype and three mutants in that background (Ryu *et al*., 2019; Jean-Baptiste *et al*., 2019; Shulse et al., 2019; Denyer *et al*., 2019). Heterogeneity of cellular responses to environmental stimuli such as high temperature and sucrose supply were also profiled in two studies (Jean-Baptiste *et al*., 2019; Shulse *et al*., 2019). These have greatly advanced our knowledge about spatiotemporal trajectories of plant root development. To maximize the usage of these public available datasets and cross-validate results from different studies, we integrated four public Arabidopsis root scRNA-seq datasets and visualized the results in our Single Cell app (Supplemental Table 4; (see data processing methods in Experimental procedures) (Jean-Baptiste *et al*., 2019; Ryu *et al*., 2019; Shulse *et al*., 2019; Denyer *et al*., 2019). By integrated analysis and visualization, we were able to compare the results of these studies and understand more about experimental variation when working with a single genotype.

We examined variation between different root single cell experiments performed on the same genotype under similar conditions. Cell-type (i.e. cluster) specific marker genes identified in one of the root experiments were used to assign cell identities to the cell clusters identified in our integrated analysis of the four experiments (Fig. 2, Supplemental Table 5; (Denyer *et al*., 2019)). Our analysis revealed that some marker genes are specific across studies – for example, pericycle/phloem (CLE26, NTL, NAT7, HCA2, FAF4). However, looking at the studies individually for the same genes indicated that there is still variation in the extent of expression (Fig. 3). FAF4 is defined as a good marker for pericycle/phloem on the basis of the data taken from Jean-Baptiste *et al*., (2019), but not based on the data from Shulse *et al*., (2019). For protoxylem, At1g14190 and UGD1 are markers based on all the studies and for trichoblasts GH9C1 and At1g07795 appear quite robust, although notably GH9C1 is also consistently found in the meristematic xylem, and undefined in the study of Shulse *et al*., (2019). When all datasets are integrated and analysed together, we observed that previously defined atrichoblast markers are in fact also found in columella, the quiescent centre (QC), cortex, meristematic xylem and trichoblast. Likewise, the cortex marker TBL41 is found in meristematic xylem, atrichoblast, trichoblast. Although atrichoblast marker genes were highly expressed in atrichoblast clusters (0, 12), they were also expressed in other cell types such as columella and trichoblast, which is consistent with a previous report (Denyer *et al*., 2019).

**Figure 2.**
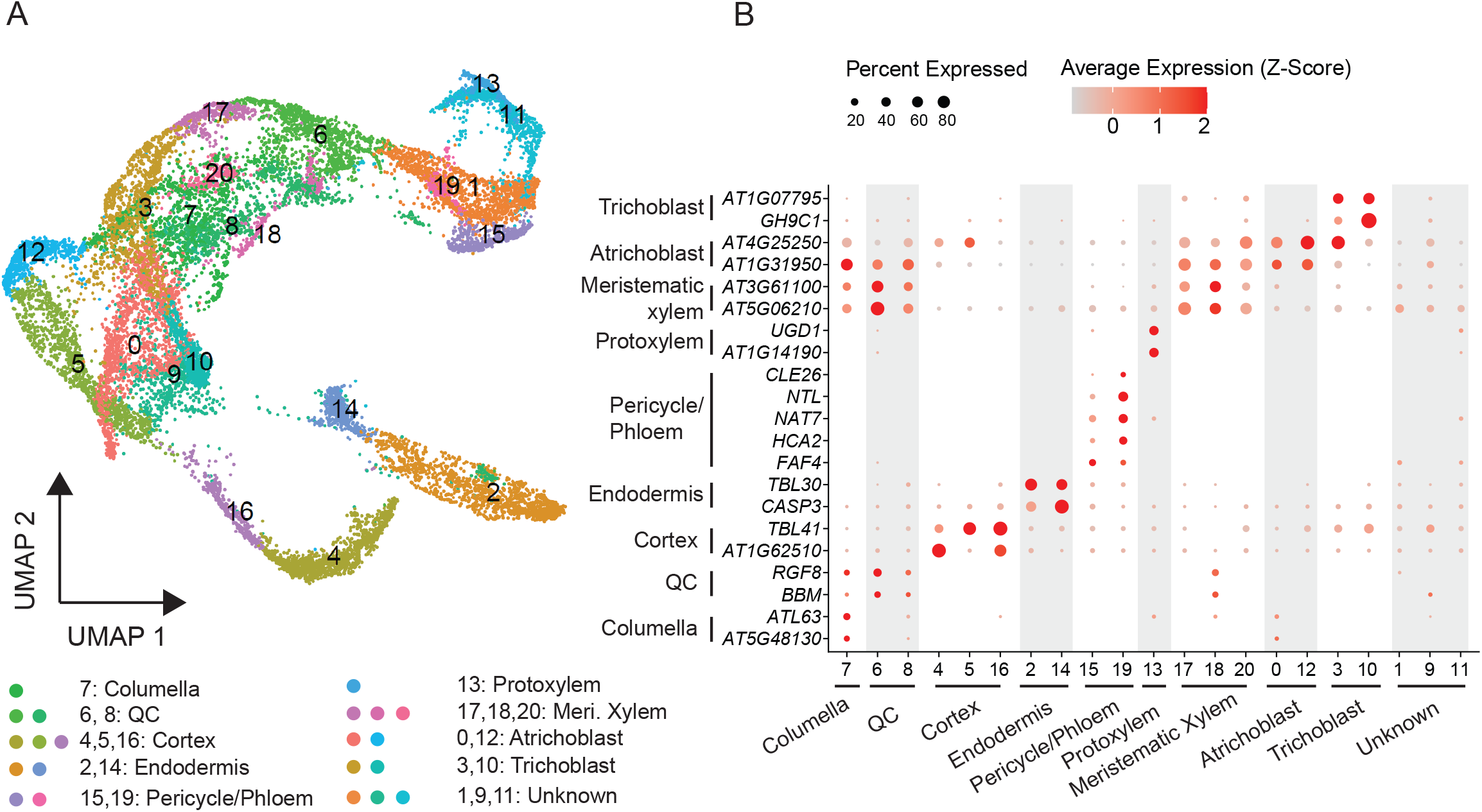
Cell Heterogeneity in Arabidopsis Root. (A) Visualization of 21 cell clusters using UMAP. Dots, individual cells; n = 16,213 cells; color, cell clusters. (B) Expression pattern of representative cluster-specific marker genes. Dot diameter, proportion of cluster cells expressing a given gene. Color, mean expression across cells in that cluster. QC: quiescent centre. The full names of selected genes are given in Supplemental Table 1.

**Figure 3.**
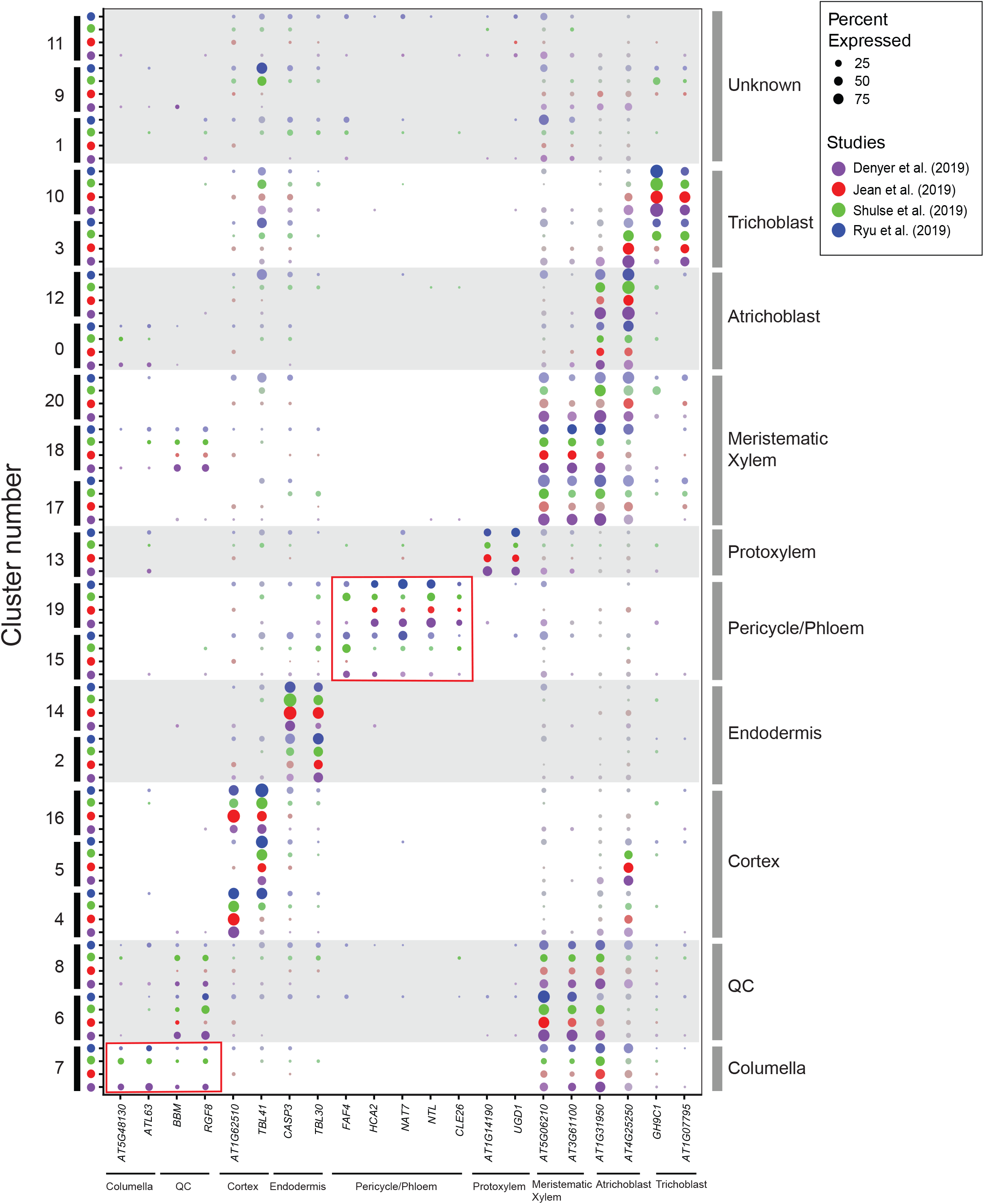
Expression of known cell-specific marker genes across four studies. Expression of cell-type marker genes in different cell clusters (left axis) and cell types (right axis) across four studies. While most cell-specific marker genes are conserved across different studies, variations of expression percentage of some marker genes were observed (red box). Dot diameter indicates the proportionof cluster cells expressing a given gene. Color indicates different studies. QC: quiescent center.

We also discovered three clusters (1,9,11) in our integrated analysis of all datasets that could not be assigned to a cell type because none of the known marker genes were expressed specifically in these clusters. These analyses reveal that while the concept of marker genes is useful for biological interpretation of individual studies, markers may vary across different experiments. This does not invalidate individual studies, rather it suggests that subtle differences in growth conditions cause some variation that become apparent when a larger set of studies is analysed. The ability to compare studies and look at them together using our app is valuable in designing future experiments and forming hypotheses. The ability to identify cell specific promoters benefits from combining many studies, but understanding variation between individual studies is also beneficial before proceeding with experimental studies.

### Mining expression profiles of phosphate transporters using the Single Cell App

Phosphate (Pi) is an important macro nutrient for plant growth and development. Pi transporters (PHTs) play a critical role in Pi uptake from the environment and translocation between organs, cell types or organelles. (Hamburger et al., 2002; Muchhal et al., 1996; Mudge et al., 2002; Shin et al., 2004). Five PHT families (PHT1 to PHT5) with different subcellular localisation have been characterized in plant based on phylogenetic analysis (Irigoyen et al., 2011; Mudge *et al*., 2002; Versaw and Harrison, 2002; Wang et al., 2017); (Zhu et al., 2012). PHOSPHATE1 (PHO1) is a Pi exporter which is responsible for loading Pi into the xylem vessels (Hamburger *et al*., 2002). In addition, 10 PHO1 homologs (PHO1;H1 to PHO1;H10) have been identified in Arabidopsis (Wang et al., 2004). While many studies have been performed to decipher the functions of the Pi transporters, a comprehensive spatial expression profile of the Pi transporters is limited (Hamburger *et al*., 2002; Khan et al., 2014; Mudge *et al*., 2002; Shin *et al*., 2004; Stefanovic et al., 2007). Thus, we examined the expression of all known genes (32 genes) from PHT and PHO1 families in the integrated Arabidopsis root dataset (Fig. 4, Supplemental Table 6). Out of the 32 genes, only 20 genes passed the quality control (see data source and processing)(Fig. 4). Consistent with a previous report (Mudge *et al*., 2002), *PHT1;1* is strongly expressed in trichoblast, where Pi is taken up from the soil (Fig. 4A, 4C, Supplemental Table 6). We also found *PHT1;1* is highly expressed in cortex, confirming an additional role of PHT1;1 in transporting Pi from epidermis into the central cylinder of the root (Karthikeyan et al., 2002) (Fig. 4A & 4CSupplemental Table 6). PHO1 was the first characterized gene in this family and is mainly expressed in stelar cells including the pericycle and xylem parenchyma cells, which is in agreement with our results (Figure 4C) (Hamburger *et al*., 2002). So far only *PHO1;H1* was shown to complement the PHO1 loss-of-function mutant, while the function of other PHO1 homologs remains unknown (Stefanovic *et al*., 2007). Seven PHO1 family genes (*PHO1, PHO1;H1, PHO1;H2, PHO1;H4, PHO1;H5, PHO1;H7, PHO1;H10*) are induced by protoplasting (Denyer *et al*., 2019), while only PHO1, PHO1;H1, PHO1;H3 and PHO1;H10 passed the quality control in our study. Interestingly, in contrast to a previous study which reported *PHO1;H3* is expressed in root vascular cylinder (Khan *et al*., 2014), we found *PHO1;H3* is mainly expressed in the endodermis (Fig. 4B, 4C). Given endodermal cells have thick cell walls restricting water or ion transport through this layer to the symplastic pathway. Therefore PHO1;H3 might also facilitate Pi transport from cortex into stelar cells through the endodermis. These significant differences between studies with respect to cell-type specific Pi transporters expression highlight the potential for comparative approaches enabled by our app to prompt further hypothesis-driven experiments.

**Figure 4.**
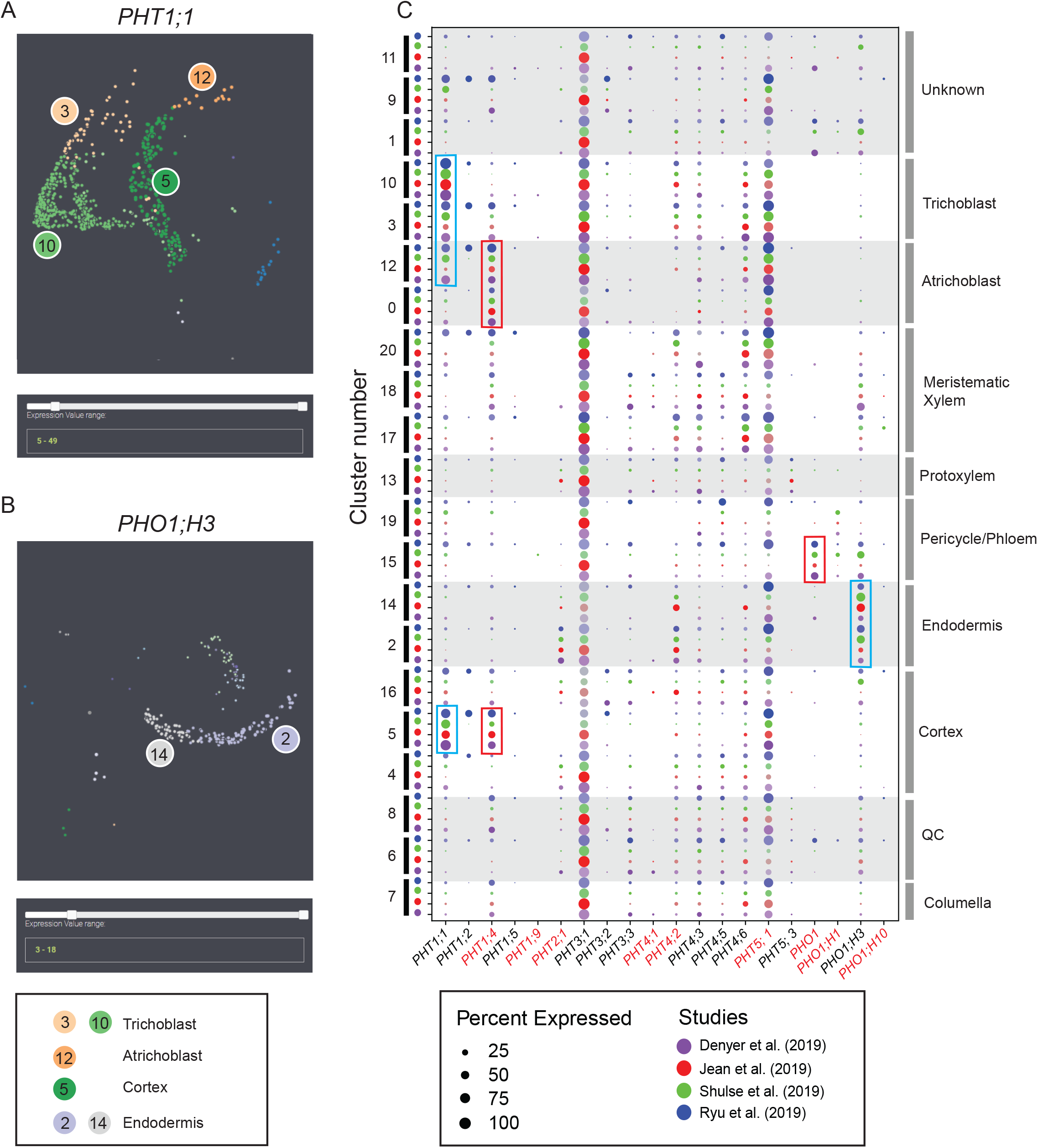
Visualization of expression of phosphate transporters in different cell types. (A) and (B), Screenshot of expression of *PHT1;1* (A) and *PHO1;H3* (B) across all studies from Single Cell App. Cells with less than 5 and 3 reads were filtered out for *PHT1;1* and *PHO1;H3*, respectively. *PHT1;1* is mainly expressed in trichoblast, atrichoblast and cortex; *PHO1;H3* is mainly expressed in endodermis. (C), Expression of expressed phosphate transpoters across four studies. Clusters highly expressing *PHT1;1* and *PHO1;H3* were highlighted with lightblue boxes. Expression percentage of *PHT1;1* in cluster 12 from Jean et al. (2019) was less than the other three studies. Protoplasting-induced genes were marked as red text. Althought induced by protoplast isolation, *PHT1;4* and *PHO1* still show cell-specific expression pattern (red boxes) with *PHT1;4* specificlly expressed in cortex and atrichoblast, *PHO1* specificly expressed in pericycle/pholem.

### Characterising single cell expression patterns of plant hormone genes in the Arabidopsis root

Plant hormones regulate plant growth, development and stress responses (Kumar, 2013; Santner et al., 2009). Essentially all plant hormones are involved in root development in some manner (Fu and Harberd, 2003; Kumar, 2013; Qin et al., 2019) (Staswick et al., 1992; Yang et al., 2017; Zhao et al., 2014; McAdam et al., 2016; Ruzicka et al., 2007; Tian et al., 2009). Hormonal signalling operates in a cell and tissue specific manner, which is critical to enable the unique functions and responses of individual cell types (Novak et al., 2017). However, there has been little transcriptomic analysis of the expression patterns of plant hormone signalling genes at the level of individual cells. The advent of single cell gene expression technologies allows us to investigate this.

To characterize the single cell expression profiles of plant hormone related genes, we examined the expression of 685 marker genes of seven hormones from previous studies; auxin (63 genes), abscisic acid (311), brassinolide (6), cytokinins (14), ethylene (3), gibberellic acid (9) and JA (279) (Birnbaum et al., 2003; Nemhauser et al., 2006; Zander et al., 2020). We computed a measure of cell-specific expression using the tissue-specificity metric tau in order to understand what proportion of hormone marker genes were expressed in specific cell-types (Yanai et al., 2005). For each gene, the average expression of all cells from the same cell type was used to calculate the expression level in this cell type. The tau value was computed using these average expression levels from different cell types. The tau metric ranges from 0 to 1, with values >0.85 indicating tissue/cluster-specific expression and values <0.15 indicative of very broad expression. One hundred and forty marker genes had tau values >0.85, with all hormones represented (Supplemental Table 7). The distribution of tau values was skewed toward the upper end of the range, centered around 0.75, indicating that most hormone marker genes were expressed in a subset of clusters rather than broadly and in the fashion of housekeeping genes (Fig. 5A).

**Figure 5.**
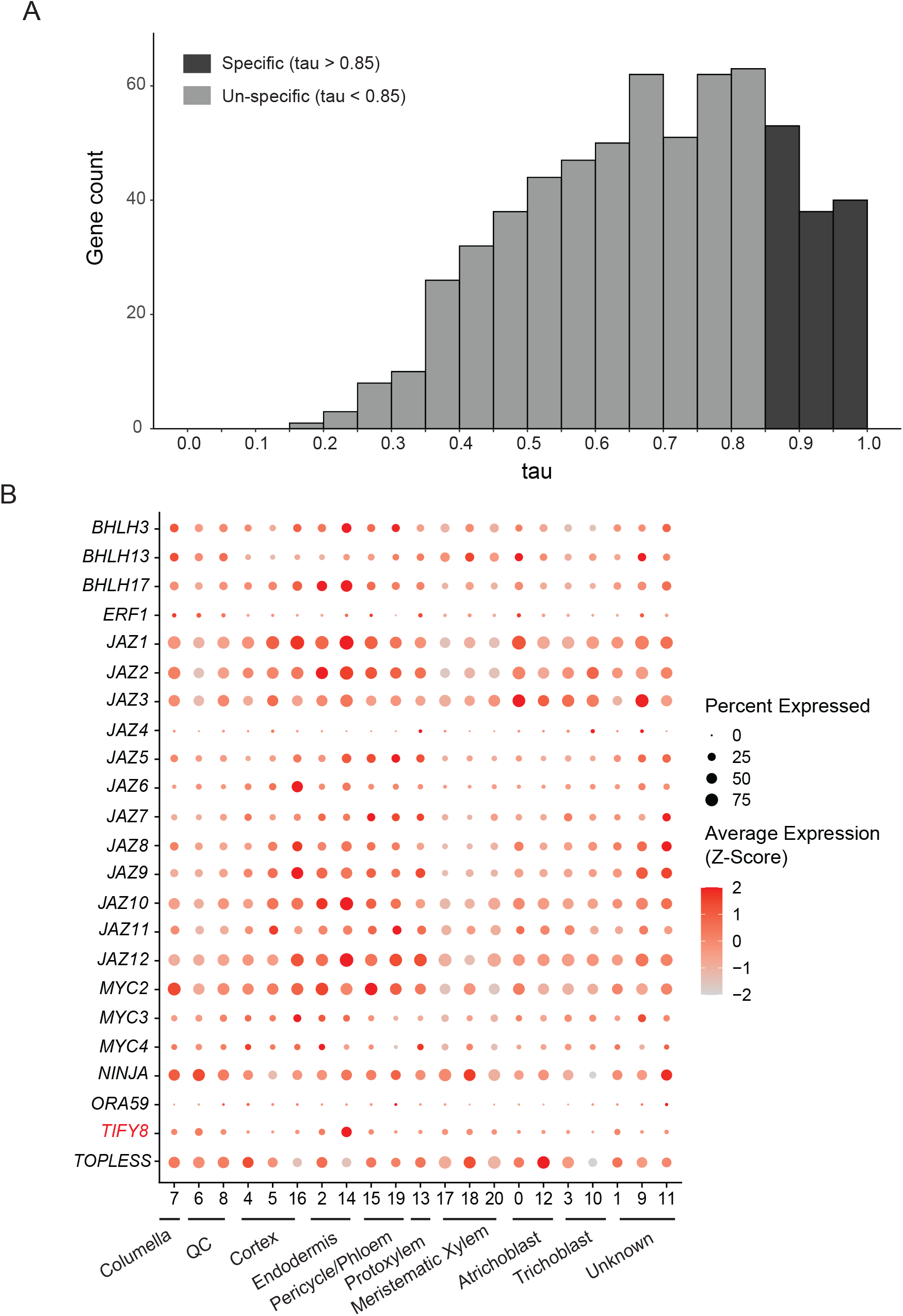
Expression of hormone marker genes in different cell types. (A) Distribution of tau values for 643 known hormone marker genes in the combined root scRNA-seq dataset. The average expression of all cells from the same cell type was calculated. This was then used to calculate a tau value across all cell clusters. Dark grey bars highlight tissue-specific values of tau > 0.85, which indicates high cluster specificity. Amongst the 643 detected genes, 132 have cell specific expression (Supplemental Table 7). (B) Expression of 23 JA marker genes in different cell types. *TIFY8* is the only of these with tau > 0.85 and is expressed specifically expressed in the endodermis.

We focused our analysis on core components of the JA transcriptional regulatory mechanism because the promoters of the *MYC2, 3, 4* transcription factors drive cell-specific reporter expression in roots (Gasperini et al., 2015). Though we observed some variation between cell-types in the scRNA-seq datasets, these transcription factors were expressed relatively broadly, suggesting differences in experimental conditions or the approaches employed may contribute to changed expression domains. We reasoned that other JA components may behave similarly and examined expression of *MYC2, 3, 4*, the *JAZ* repressors and other related factors (Fig. 5B). *NINJA* was expressed broadly across root cell-types, consistent with the previously reported behaviour of its promoter (Gasperini *et al*., 2015). We observed that only one component, *TIFY8*, exceeded the tau threshold of 0.85, which indicated cluster-specific expression in the endodermis cluster 14. TIFY8 interacts with proteins that regulate root meristem initiation and this expression pattern perhaps reflects the participation of the endodermis in lateral root formation (Cuellar Perez et al., 2014; Torres-Martinez et al., 2019). There was, however, clear variation across clusters in both expression level and proportion of cells expressing other JA transcriptional regulators for those components which did not exceed the tau threshold. For example, expression of *JAZ* repressors was generally lower in columella, QC and meristematic xylem clusters. Expression of *JAZ5* was highest in endodermis, pericycle/phloem and protoxylem clusters, whilst *JAZ6* expression was highest in cortex cluster 16. These results likely indicate that *JAZ* repressors have cell-type specific functions in gene expression regulation in the root, potentially explaining why the *JAZ/TIFY* family is relatively large.

## Discussion

scRNA-seq has been used by plant scientists since 2013 and related publications have dramatically increased recently due to the advent of droplet-based scRNA-seq technologies (Chen et al., 2021). While the increasing number of scRNA-seq studies has advanced our knowledge about plant cell heterogeneity, mining public data is challenging for researchers with limited computational expertise. In this study, we developed Single Cell App, a web-based platform for scRNA-seq data visualization. Unlike other online scRNA-seq tools which focus on data processing, our Single Cell App provides a user-friendly data visualization interface (Feng *et al*., 2019; Franzen and Bjorkegren, 2020; Mädler et al., 2020). In addition to a common gene-based search function, Single Cell App also provides a function-based search option which is useful for users to examine expression of genes related to a specific biological process. We also presented an integrated Arabidopsis root dataset from four publications (Ryu *et al*., 2019; Shulse *et al*., 2019; Jean-Baptiste *et al*., 2019; Denyer *et al*., 2019), showing the utility of the Single Cell App to define hypotheses for experimental testing. Together with the Single Cell App, public datasets provide a powerful resource for researchers who wish to examine gene expression in a single cell resolution.

In our study, we annotated cell clusters from an integrated, multi-experiment root scRNA-seq dataset using well-characterized cell marker genes. We identified similar patterns of cell identity to previous studies (Fig. 2) (Ryu *et al*., 2019; Shulse *et al*., 2019; Jean-Baptiste *et al*., 2019; Denyer *et al*., 2019). Unexpectedly, we found that the specificity of some marker genes varied between studies, with some genes expressed in different cell types compared to previous publications. *PHO1;H3* was reported to specifically express in stele while it is mainly expressed in endodermis in this study (Fig. 4B, 4C), (Khan *et al*., 2014). These differences may result from different environmental stimuli or growth stages between studies. For example, stele-specific expression of *PHO1;H3* was detected in plants grown in an zinc deficient environment while data analysed here was obtained from seedlings supplied with sufficient nutrients (Khan *et al*., 2014). Indeed, cell heterogeneity in response to environmental stimuli has been previously reported (Berkowitz *et al*., 2021; Jean-Baptiste *et al*., 2019; Shulse *et al*., 2019). We also observed here that a variety of genes encoding phosphate transporters were induced by protoplasting, which would normally be discarded in analysis. However, these genes were still expressed in a cell specific manner (Fig. 4C). Likewise with the analysis of hormone response genes it demonstrated that many genes are expressed in a cell enriched manner, indicating that data from whole organ studies that make interpretation of signalling and functional pathways, may need to be re-examined as all the genes are not expressed in the same cell. Thus, these ‘pathways’ may not exist in any one cell suggesting heterogeneity in cell transcriptomes. Quantitative imaging of single cell transcriptional dynamics in plant cells have demonstrated that there are large differences between neighbouring cells, that is consistent with the conclusion that we have reached here with the comparison cell specific transcriptomes (Alamos et al., 2021; Hani et al., 2021).

In conclusion due to differences in growth conditions and cell heterogeneity in transcriptional dynamics it is important that different studies looking at cell specific transcriptomes can be readily compared. This will allow a synergistic use of the data and prompt hypothesis and discovery-based experiments that will give greater insight into cell specific function and heterogeneity.

## Methods

### Single cell app architecture

The architecture of the Single Cell app uses the Microsoft Azure public clouds (Supplemental Figure 1). It is primarily based on Platform as a Service (PaaS) and Software as a Service (SaaS) solution. This provides automatic scalability, such that when additional storage or compute resources are required to perform an analysis or visualisation, the cloud backend will automatically make those resources available to the app, within a set of specified parameters. It also allows easy deployment *via* the Loome platform. Users connect to the Single Cell app over the internet using a web browser, after typing a custom web address (URL) that can be associated with an institutional domain. The web interface for the Single Cell application presents the users with two main areas: 1) Upload Experiment: provides the functionality to upload an expression matrix, a coordinates files, a GO slim file, and metadata associated with the data so that it can be identified as part of an experiment. 2) Experiment Analysis: provides the visualisation of the data for experiments that have been previously uploaded, with rich controls to search, filter and modify the data that is being visualised. The user can also export the data being visualised, in GO slim format or comma-separated values (CSV). When a user uploads new experiment data, this data is processed by the Loome Integrate agent and stored in the Application and Visualisation Databases. The agent runs in a small Docker container instance, and uses cloud blob storage to maintain logs and state. All the application components are contained within a resource group, which is a container that holds related resources for an Azure solution. This resource group also provides consolidated cost analysis and the ability to assign budgets and alerts, based on usage.

**Table 1.** The specifications for components are shown.

| **Component** | **Type** | **Specifications** |
| --- | --- | --- |
| **App Services** | App Service Plan | Auto-scaling from 1 to 3 instances, based on CPU utilisation. S1 pricing tier. |
| **Visualisation Web app** | App Service | Managed by App Services. Includes custom domain for the Single Cell App. |
| **File Uploader Web app** | App Service | Managed by App Services. |
| **SQL Server** | SQL Server | Logical SQL server. Automatic tuning and transparent data encryption. |
| **Application Database** | SQL Database | Serverless, Gen5, 16 vCores |
| **Visualisation Database** | SQL Database | General Purpose. S1 pricing tier. |
| **Loome Integrate agent** | Azure Container Instance | Linux, 1 instance, 2 vCores, 8 GiB memory |
| **Storage blob** | Storage account | General Purpose v1 |

### Data sources and processing

To prepare the root single cell data for visualization we downloaded the raw count data from the respective GEO repositories (Supplemental Table 4). Note that TAIR10 genome were used to generate the raw count data across the four public data by the authors (Ryu *et al*., 2019; Shulse *et al*., 2019; Jean-Baptiste *et al*., 2019; Denyer *et al*., 2019). It is first necessary to process the raw data and best practice for doing so has been well-reviewed elsewhere (Shaw *et al*., 2021). Our approach was to use Seurat for down-stream data processing, including quality control, data normalization, identification of highly variable features, dimensional reduction and cell clustering (Version 3.2.2) (Stuart et al., 2019). In brief, to discard low quality cells, cells expressing less than 800 genes or 3000 unique molecule identifiers were filtered out. Cells with more than 20% mitochondrial sequences were also removed to exclude dead cells. Genes expressed in less than 30 cells were also dismissed. After filtering, 21557 expressed genes across 16213 cells were retained. The raw count data were normalized using SCTransform and different datasets were integrated using Seurat (Version 3.2.2, (Stuart *et al*., 2019; Hafemeister and Satija, 2019)). Dimensionality reduction was performed on the normalized data using principle component analysis followed by Uniform Manifold Approximation and Projection (UMAP) to visualize the data structure. Cell clusters were characterized using the function ‘FindClusters’ from Seurat package with ‘‘resolution = 0.6’’, resulting 21 cell clusters (Fig. 2). Gene expression is affected by the protoplast isolation that is used during plant single cell sample preparation, which might bias cell clustering. To remove this potential bias, we excluded protoplast-induced genes identified by Denyer *et al*., (2019) in dimensionality reduction and cell clustering step. The expression matrix, UMAP with three components and the GO annotation of Arabidopsis were uploaded into the Single Cell App for the interactive data exploration.

### Calculation of cell/cluster specificity index (tau)

The metric tau was used as an index of cluster-specific expression. It was computed as previously described (Yanai *et al*., 2005). The average expression of an individual gene was calculated across all cells from the same cell type. The tau value was then computed using these average expression levels from the different cell types.

## Author contributions

M.G.L. and J.W. conceived the project and C.Y. and O.B. carried out the analysis of the biological datasets. F.A. and M.B. designed and implemented the cloud-based architecture for uploading, processing and visualising the data. J.W. and F.A. drafted the manuscript with extensive editing by all authors.

## Acknowledgments

F.A. and M.B. are employed by BizData who host the Application on the Loome platform as a fee for service product. M.G.L. and J.W. are funded by the Australian Research Council Industrial transformation Research Hub for Medicinal Agriculture (IH180100006).

## Figure legends

**Figure 1. Screenshot of the Single Cell Browser Layout**.

(A) Meta information of the selected experiment. Clicking the “View Experiment Details” will provide more information about the experiment.

(B) Color mode of displaying data and filtering cells based on gene expression level. In Cluster mode, each color represents a cell cluster while in Values mode, the color represents the expression level of a searched gene. Cells can be filtered by dragging the Expression Value range bar.

(C) Search tab. Genes can be searched based on the Gene Ontology or gene id.

(D) Three dimensions of the UMAP clustering. Each dot represents a cell.

(E) Check box for dataset selection, this can be used for showing cells from specified dataset.

(F) Cluster legend. Each color represents a cluster, the number shows the cluster id. Clicking each cluster will hide/show the cells from this cluster.

**Figure 2. Cell Heterogeneity in Arabidopsis Root**.

(A) Visualization of 21 cell clusters using UMAP. Dots, individual cells; n = 16,213 cells; color, cell clusters.

(B) Expression pattern of representative cluster-specific marker genes. Dot diameter, proportion of cluster cells expressing a given gene. Color, mean expression across cells in that cluster. QC: quiescent centre. The full names of selected genes are given in Supplemental Table 1

**Figure 3. Expression of known cell-specific marker genes across four studies**.

Expression of cell-type marker genes in different cell clusters (left axis) and cell types (right axis) across four studies. While most cell-specific marker genes are conserved across different studies, variations of expression percentage of some marker genes were observed (red box). Dot diameter indicates the proportion of cluster cells expressing a given gene. Color indicates different studies. QC: quiescent center.

**Figure 4. Visualization of expression of phosphate transporters in different cell types**.

(A) and (B), Single Cell App screenshot of for expression *PHT1;1* (A) and *PHO1;H3* (B) across used studies Cells with less than 5 and 3 reads were filtered out for *PHT1;1* and *PHO1;H3*, respectively. *PHT1;1* is mainly expressed in trichoblast, atrichoblast and cortex; *PHO1;H3* is mainly expressed in endodermis.

(C), Expression of expressed phosphate transporters across the four indicated studies. Clusters of cells highly expressing *PHT1;1* and *PHO1;H3* are represented by light blue boxes. Expression percentage of *PHT1;1* in cluster 12 from Jean et al. (2019) was less than the other three studies. Protoplasting-induced genes are highlighted in red text. Although induced by protoplast isolation, *PHT1;4* and *PHO1* are still expressed cell-specifically (red boxes), with *PHT1;4* expressed in cortex and atrichoblast, and *PHO1* expressed in pericycle/phloem.

**Figure 5. Expression of hormone marker genes in different cell types**.

(A) Distribution of tau values for 643 known hormone marker genes in the combined root scRNA-seq dataset. The average expression of all cells from the same cell type was calculated. This was then used to calculate a tau value across all cell clusters. Dark grey bars highlight tissue-specific values of tau > 0.85, which indicates high cluster specificity. Amongst the 643 detected genes, 132 have cell specific expression (Supplemental Table 7).

(B) Expression of 23 JA marker genes in different cell types. *TIFY8* is the only of these with tau > 0.85 and is expressed specifically expressed in the endodermis.

---

**Supplemental Figure 1.**
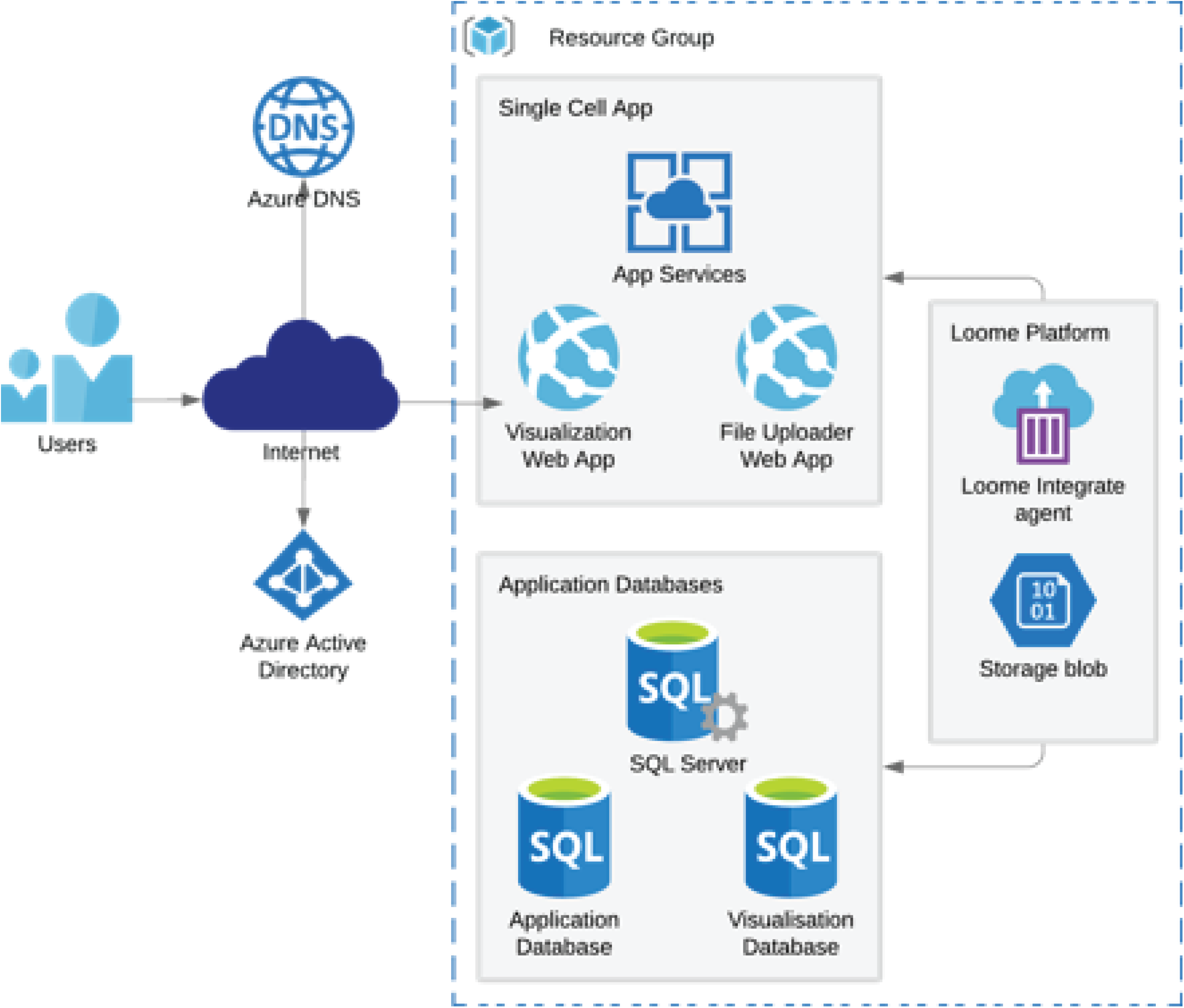
The architecture of the Single Cell app.

## References

Alamos, S., Reimer, A., Niyogi, K.K., and Garcia, H.G. (2021). Quantitative imaging of RNA polymerase II activity in plants reveals the single-cell basis of tissue-wide transcriptional dynamics. Nat Plants 7:1037–1049. 10.1038/s41477-021-00976-0.

Berkowitz, O., Xu, Y., Liew, L.C., Wang, Y., Zhu, Y., Hurgobin, B., Lewsey, M.G., and Whelan, J. (2021). RNA-seq analysis of laser microdissected Arabidopsis thaliana leaf epidermis, mesophyll and vasculature defines tissue-specific transcriptional responses to multiple stress treatments. Plant J 107:938–955 10.1111/tpj.15314.

Birnbaum, K., Shasha, D.E., Wang, J.Y., Jung, J.W., Lambert, G.M., Galbraith, D.W., and Benfey, P.N. (2003). A gene expression map of the Arabidopsis root. Science 302:1956–1960. 10.1126/science.1090022.

Chen, H., Yin, X., Guo, L., Yao, J., Ding, Y., Xu, X., Liu, L., Zhu, Q.H., Chu, Q., and Fan, L. (2021). PlantscRNAdb: A database for plant single-cell RNA analysis. Mol Plant 14:855–857. 10.1016/j.molp.2021.05.002.

Cuellar Perez, A., Nagels Durand, A., Vanden Bossche, R., De Clercq, R., Persiau, G., Van Wees, S.C., Pieterse, C.M., Gevaert, K., De Jaeger, G., Goossens, A., et al. (2014). The non-JAZ TIFY protein TIFY8 from Arabidopsis thaliana is a transcriptional repressor. PLoS One 9:e84891. 10.1371/journal.pone.0084891.

Denyer, T., Ma, X., Klesen, S., Scacchi, E., Nieselt, K., and Timmermans, M.C.P. (2019). Spatiotemporal Developmental Trajectories in the Arabidopsis Root Revealed Using High-Throughput Single-Cell RNA Sequencing. Dev Cell 48:840–852 e845. 10.1016/j.devcel.2019.02.022.

Drapek, C., Sparks, E.E., and Benfey, P.N. (2017). Uncovering Gene Regulatory Networks Controlling Plant Cell Differentiation. Trends Genet 33:529–539. 10.1016/j.tig.2017.05.002.

Feng, D., Whitehurst, C.E., Shan, D., Hill, J.D., and Yue, Y.G. (2019). Single Cell Explorer, collaboration-driven tools to leverage large-scale single cell RNA-seq data. BMC Genomics 20:676. 10.1186/s12864-019-6053-y.

Franzen, O., and Bjorkegren, J.L.M. (2020). alona: a web server for single-cell RNA-seq analysis. Bioinformatics 36:3910–3912. 10.1093/bioinformatics/btaa269.

Fu, X., and Harberd, N.P. (2003). Auxin promotes Arabidopsis root growth by modulating gibberellin response. Nature 421:740–743. 10.1038/nature01387.

Gasperini, D., Chetelat, A., Acosta, I.F., Goossens, J., Pauwels, L., Goossens, A., Dreos, R., Alfonso, E., and Farmer, E.E. (2015). Multilayered Organization of Jasmonate Signalling in the Regulation of Root Growth. PLoS Genet 11:e1005300. 10.1371/journal.pgen.1005300.

Hafemeister, C., and Satija, R. (2019). Normalization and variance stabilization of single-cell RNA-seq data using regularized negative binomial regression. Genome Biol 20:296. 10.1186/s13059-019-1874-1.

Hamburger, D., Rezzonico, E., MacDonald-Comber Petetot, J., Somerville, C., and Poirier, Y. (2002). Identification and characterization of the Arabidopsis PHO1 gene involved in phosphate loading to the xylem. Plant Cell 14:889–902. 10.1105/tpc.000745.

Hani, S., Cuyas, L., David, P., Secco, D., Whelan, J., Thibaud, M.C., Merret, R., Mueller, F., Pochon, N., Javot, H., et al. (2021). Live single-cell transcriptional dynamics via RNA labelling during the phosphate response in plants. Nat Plants 7:1050–1064. 10.1038/s41477-021-00981-3.

Irigoyen, S., Karlsson, P.M., Kuruvilla, J., Spetea, C., and Versaw, W.K. (2011). The sink-specific plastidic phosphate transporter PHT4;2 influences starch accumulation and leaf size in Arabidopsis. Plant Physiol 157:1765–1777. 10.1104/pp.111.181925.

Jean-Baptiste, K., McFaline-Figueroa, J.L., Alexandre, C.M., Dorrity, M.W., Saunders, L., Bubb, K.L., Trapnell, C., Fields, S., Queitsch, C., and Cuperus, J.T. (2019). Dynamics of Gene Expression in Single Root Cells of Arabidopsis thaliana. Plant Cell 31:993–1011. 10.1105/tpc.18.00785.

Karthikeyan, A.S., Varadarajan, D.K., Mukatira, U.T., D’Urzo, M.P., Damsz, B., and Raghothama, K.G. (2002). Regulated expression of Arabidopsis phosphate transporters. Plant Physiol 130:221–233. 10.1104/pp.020007.

Khan, G.A., Bouraine, S., Wege, S., Li, Y., de Carbonnel, M., Berthomieu, P., Poirier, Y., and Rouached, H. (2014). Coordination between zinc and phosphate homeostasis involves the transcription factor PHR1, the phosphate exporter PHO1, and its homologue PHO1;H3 in Arabidopsis. J Exp Bot 65:871–884. 10.1093/jxb/ert444.

Kumar, P.P. (2013). Regulation of biotic and abiotic stress responses by plant hormones. Plant Cell Rep 32:943. 10.1007/s00299-013-1460-z.

Liu, Q., Liang, Z., Feng, D., Jiang, S., Wang, Y., Du, Z., Li, R., Hu, G., Zhang, P., Ma, Y., et al. (2021). Transcriptional landscape of rice roots at the single-cell resolution. Mol Plant 14:384–394. 10.1016/j.molp.2020.12.014.

Mädler, S.C., Julien-Laferriere, A., Wyss, L., Phan, M., Kang, A.S.W., Ulrich, E., Schmucki, R., Zhang, J.D., Ebeling, M., Badi, L., et al. (2020). Besca, a single-cell transcriptomics analysis toolkit to accelerate translational research. BioRxiv https://doi.org/10.1101/2020.08.11.245795.

McAdam, S.A., Brodribb, T.J., and Ross, J.J. (2016). Shoot-derived abscisic acid promotes root growth. Plant Cell Environ 39:652–659. 10.1111/pce.12669.

Moussa, M., and Mandoiu, II (2021). SC1: A Tool for Interactive Web-Based Single-Cell RNA-Seq Data Analysis. J Comput Biol 28:820–841. 10.1089/cmb.2021.0051.

Muchhal, U.S., Pardo, J.M., and Raghothama, K.G. (1996). Phosphate transporters from the higher plant Arabidopsis thaliana. Proc Natl Acad Sci U S A 93:10519–10523. 10.1073/pnas.93.19.10519.

Mudge, S.R., Rae, A.L., Diatloff, E., and Smith, F.W. (2002). Expression analysis suggests novel roles for members of the Pht1 family of phosphate transporters in Arabidopsis. Plant J 31:341–353. 10.1046/j.1365-313x.2002.01356.x.

Nemhauser, J.L., Hong, F., and Chory, J. (2006). Different plant hormones regulate similar processes through largely nonoverlapping transcriptional responses. Cell 126:467–475. 10.1016/j.cell.2006.05.050.

Novak, O., Napier, R., and Ljung, K. (2017). Zooming In on Plant Hormone Analysis: Tissue-and Cell-Specific Approaches. Annu Rev Plant Biol 68:323–348. 10.1146/annurev-arplant-042916-040812.

Qin, H., He, L., and Huang, R. (2019). The Coordination of Ethylene and Other Hormones in Primary Root Development. Front Plant Sci 10:874. 10.3389/fpls.2019.00874.

Ruzicka, K., Ljung, K., Vanneste, S., Podhorska, R., Beeckman, T., Friml, J., and Benkova, E. (2007). Ethylene regulates root growth through effects on auxin biosynthesis and transport-dependent auxin distribution. Plant Cell 19:2197–2212. 10.1105/tpc.107.052126.

Ryu, K.H., Huang, L., Kang, H.M., and Schiefelbein, J. (2019). Single-Cell RNA Sequencing Resolves Molecular Relationships Among Individual Plant Cells. Plant Physiol 179:1444–1456. 10.1104/pp.18.01482.

Santner, A., Calderon-Villalobos, L.I., and Estelle, M. (2009). Plant hormones are versatile chemical regulators of plant growth. Nat Chem Biol 5:301–307. 10.1038/nchembio.165.

Shahan, R., Nolan, T.M., and Benfey, P.N. (2021). Single-cell analysis of cell identity in the Arabidopsis root apical meristem: insights and opportunities. J Exp Bot erab228 10.1093/jxb/erab228.

Shaw, R., Tian, X., and Xu, J. (2021). Single-Cell Transcriptome Analysis in Plants: Advances and Challenges. Mol Plant 14:115–126. 10.1016/j.molp.2020.10.012.

Shin, H., Shin, H.S., Dewbre, G.R., and Harrison, M.J. (2004). Phosphate transport in Arabidopsis: Pht1;1 and Pht1;4 play a major role in phosphate acquisition from both low- and high-phosphate environments. Plant J 39:629–642. 10.1111/j.1365-313X.2004.02161.x.

Shulse, C.N., Cole, B.J., Ciobanu, D., Lin, J., Yoshinaga, Y., Gouran, M., Turco, G.M., Zhu, Y., O’Malley, R.C., Brady, S.M., et al. (2019). High-Throughput Single-Cell Transcriptome Profiling of Plant Cell Types. Cell Rep 27:2241–2247 e2244. 10.1016/j.celrep.2019.04.054.

Staswick, P.E., Su, W., and Howell, S.H. (1992). Methyl jasmonate inhibition of root growth and induction of a leaf protein are decreased in an Arabidopsis thaliana mutant. Proc Natl Acad Sci U S A 89:6837–6840. 10.1073/pnas.89.15.6837.

Stefanovic, A., Ribot, C., Rouached, H., Wang, Y., Chong, J., Belbahri, L., Delessert, S., and Poirier, Y. (2007). Members of the PHO1 gene family show limited functional redundancy in phosphate transfer to the shoot, and are regulated by phosphate deficiency via distinct pathways. Plant J 50:982–994. 10.1111/j.1365-313X.2007.03108.x.

Stuart, T., Butler, A., Hoffman, P., Hafemeister, C., Papalexi, E., Mauck, W.M., 3rd, Hao, Y., Stoeckius, M., Smibert, P., and Satija, R. (2019). Comprehensive Integration of Single-Cell Data. Cell 177:1888–1902 e1821. 10.1016/j.cell.2019.05.031.

Tian, Q.Y., Sun, P., and Zhang, W.H. (2009). Ethylene is involved in nitrate-dependent root growth and branching in Arabidopsis thaliana. New Phytol 184:918–931. 10.1111/j.1469-8137.2009.03004.x.

Torres-Martinez, H.H., Rodriguez-Alonso, G., Shishkova, S., and Dubrovsky, J.G. (2019). Lateral Root Primordium Morphogenesis in Angiosperms. Front Plant Sci 10:206. 10.3389/fpls.2019.00206.

Toufighi, K., Brady, S.M., Austin, R., Ly, E., and Provart, N.J. (2005). The Botany Array Resource: e-Northerns, Expression Angling, and promoter analyses. Plant J 43:153–163. 10.1111/j.1365-313X.2005.02437.x.

Versaw, W.K., and Harrison, M.J. (2002). A chloroplast phosphate transporter, PHT2;1, influences allocation of phosphate within the plant and phosphate-starvation responses. Plant Cell 14:1751–1766. 10.1105/tpc.002220.

Wang, D., Lv, S., Jiang, P., and Li, Y. (2017). Roles, Regulation, and Agricultural Application of Plant Phosphate Transporters. Front Plant Sci 8:817. 10.3389/fpls.2017.00817.

Wang, Y., Ribot, C., Rezzonico, E., and Poirier, Y. (2004). Structure and expression profile of the Arabidopsis PHO1 gene family indicates a broad role in inorganic phosphate homeostasis. Plant Physiol 135:400–411. 10.1104/pp.103.037945.

Xu, X., Crow, M., Rice, B.R., Li, F., Harris, B., Liu, L., Demesa-Arevalo, E., Lu, Z., Wang, L., Fox, N., et al. (2021). Single-cell RNA sequencing of developing maize ears facilitates functional analysis and trait candidate gene discovery. Dev Cell 56:557–568 e556. 10.1016/j.devcel.2020.12.015.

Yanai, I., Benjamin, H., Shmoish, M., Chalifa-Caspi, V., Shklar, M., Ophir, R., Bar-Even, A., Horn-Saban, S., Safran, M., Domany, E., et al. (2005). Genome-wide midrange transcription profiles reveal expression level relationships in human tissue specification. Bioinformatics 21:650–659. 10.1093/bioinformatics/bti042.

Yang, Z.B., He, C., Ma, Y., Herde, M., and Ding, Z. (2017). Jasmonic Acid Enhances Al-Induced Root Growth Inhibition. Plant Physiol 173:1420–1433. 10.1104/pp.16.01756.

Zander, M., Lewsey, M.G., Clark, N.M., Yin, L., Bartlett, A., Saldierna Guzman, J.P., Hann, E., Langford, A.E., Jow, B., Wise, A., et al. (2020). Integrated multi-omics framework of the plant response to jasmonic acid. Nat Plants 6:290–302. 10.1038/s41477-020-0605-7.

Zhang, H., Zhang, F., Yu, Y., Feng, L., Jia, J., Liu, B., Li, B., Guo, H., and Zhai, J. (2020). A Comprehensive Online Database for Exploring approximately 20,000 Public Arabidopsis RNA-Seq Libraries. Mol Plant 13:1231–1233. 10.1016/j.molp.2020.08.001.

Zhang, T.Q., Chen, Y., Liu, Y., Lin, W.H., and Wang, J.W. (2021). Single-cell transcriptome atlas and chromatin accessibility landscape reveal differentiation trajectories in the rice root. Nat Commun 12:2053. 10.1038/s41467-021-22352-4.

Zhao, Y., Xing, L., Wang, X., Hou, Y.J., Gao, J., Wang, P., Duan, C.G., Zhu, X., and Zhu, J.K. (2014). The ABA receptor PYL8 promotes lateral root growth by enhancing MYB77-dependent transcription of auxin-responsive genes. Sci Signal 7:ra53. 10.1126/scisignal.2005051.

Zhu, W., Miao, Q., Sun, D., Yang, G., Wu, C., Huang, J., and Zheng, C. (2012). The mitochondrial phosphate transporters modulate plant responses to salt stress via affecting ATP and gibberellin metabolism in Arabidopsis thaliana. PLoS One 7:e43530. 10.1371/journal.pone.0043530.

Zimmermann, P., Hirsch-Hoffmann, M., Hennig, L., and Gruissem, W. (2004). GENEVESTIGATOR. Arabidopsis microarray database and analysis toolbox. Plant Physiol 136:2621–2632. 10.1104/pp.104.046367.

